# Prefrontal and ventral striatal dendritic morphology: effects of life-long complex housing and amphetamine administration

**DOI:** 10.1101/2024.07.08.602601

**Authors:** Bryan Kolb, Terry E. Robinson

## Abstract

Complex housing is one of the most effective experiences in producing plastic changes in the brain. For example, animals living in complex environments show widespread synaptic changes both in cerebral cortex and the striatum. Similarly, repeatedly treating animals with psychomotor stimulants such as amphetamine also induces changes in prefrontal cortex and the striatum. The purpose of the current study was to determine the effects of life-long housing in complex environments versus standard laboratory caging and this experience influenced the later effects of amphetamine. Both male and female Long-Evans rats were placed in complex environments for about 110 days, beginning at conception, until adulthood at which time they were administered saline or amphetamine daily (1 mg/kg, IP) for 14 days. A week later the brains were harvested and processed for Golgi-Cox staining to analyze dendritic length, branching, and spine density in prefrontal cortex (areas Cg3 and AID) and Nucleus Accumbens (NAcc). The results showed that the prolonged period of enriched housing produced significant synaptic changes in all three measures in all three areas measured, but the effects differed in the two sexes. Amphetamine produced large synaptic changes in Cg3 and NAcc in males but only spine changes in those regions in females. Complex housing did not interact with the later effects of amphetamine administration. Thus, both complex housing and amphetamine can produce a range of synaptic changes depending upon sex and area examined. Furthermore, the effect of complex housing varies depending on the details of when complex housing is begun and how long it lasts.

---

Relative to lab animals housed in standard cages, those housed in a relatively complex (“enriched”) environment show alterations in neuronal morphology in many cortical regions, including increased cortical thickness and alterations in cellular morphology, such as increased dendritic length, dendritic complexity, spine density, and synaptic size (e.g., Galaj et al., 2020; Juraska, 1990; Kolb et al., 2003a; Rosenzweig & Bennet, 1996; Sirevaag & Greenough, 1988; Smail et al., 2020). Training in a range of motor and cognitive tasks produce similar changes in specific cortical regions (e.g., Greenough & Chang, 1989; Kolb et al., 2008; Whithers & Greenough, 1989). Most previous studies focused on sensory/motor regions and the hippocampus, but one study reported large enrichment-related increases in dendritic length and complexity on pyramidal cells in the parietal cortex and medium spiny neurons in the nucleus accumbens (NAcc), but no effects on these measures in the medial frontal cortex (mPFC) (Kolb et al., 2003b). However, spine density was increased in all three regions (see also Ashokan et al., 2018).

Prefrontal neurons are very plastic in response to other experiences (Forgie & Kolb, 2003; Robinson & Kolb, 1997; Robinson & Kolb, 2004; Tian et al., 2010; Kolb et a., 1997; Comeau et al, 2010), so this does not account for the small effects of complex housing on mPFC neurons discussed above. However, our earlier report has two shortcomings that may account for the small effects. One, only female rats were used and, two, only one prefrontal region (mPFC) was studied. Given there are sex differences in neuronal morphology in prefrontal regions (e.g., Kolb & Stewart, 1991; Willing & Juraska, 2015) and many experiences differentially affect mPFC (Zilles area Cg3) and orbital regions (Zilles area AID) (e.g., Bell et al., 2010; Robinson & Kolb, 2004), one goal of the present study was to replicate our original findings, but to now include both female and male rats and to compare the effects of complex housing in both the mPFC and orbital frontal cortex. To maximize the effects of complex housing one group of animals lived in the complex environment their entire lives (see Sampedro-Piquero & Begega, 2017) while another group were group-housed in standard laboratory cages.

In addition to complex housing, treatment with a number of potentially addictive drugs, including amphetamine, cocaine, or nicotine, produce marked effects on dendritic morphology in the mPFC, other cortical areas and the nucleus accumbens (NAcc) (Robinson and Kolb, 2004; Hamilton and Kolb, 2005), and these drug-induced effects interact with the effects of complex housing. For example, treatment with amphetamine, cocaine, or nicotine, using a regimen that produces behavioral sensitization, attenuates the subsequent effects of complex housing on neurons in NAcc and parietal cortex (Hamilton & Kolb, 2005; Kolb et al., 2003c). However, the reverse may not be true because Hamilton & Kolb (2005) found no effect of complex housing on the ability of subsequent treatment with nicotine to induce alterations in dendritic morphology.

Given that it has been suggested that environmental enrichment may have therapeutic effects on substance use disorders (e.g., Galaj et al., 2020) we thought this question deserves further examination. Thus, in the current study we housed rats in a complex environment from conception and then after 90 postnatal days quantified the ability of amphetamine to induce behavioral sensitization and alter dendritic morphology of cells in the mPFC (Cg3), orbital frontal cortex (AID) and nucleus accumbens. The importance of beginning enrichment so early is reinforced by a study showing that the milk from rat dams in enriched environments is altered significantly including greater triglyceride levels and microbial diversity as well as large changes in gene expression (DeRosa et al., 2022). These differences are associated with heavier offspring body weight and increased sociability. It therefore would not be surprising to find differences in a wide range of behaviors, including response to psychoactive drugs in the complex-housed animals.

## Materials and methods

### Animals

Long-Evans rats (F=32; M=32) derived from Charles-Rivers strains, from 8 different litters, were used. Four litters were born and raised in a complex housing environment and four litters were born and raised in Plexiglas cages (see below). The animals’ sex was determined at weaning and the two sexes were segregated into female or male groups with eight animals per group in the complex environments and two per group in the control caging. At 90 days of age the animals were randomly divided into groups getting amphetamine or saline. Three animals died by 90 days leaving the following n’s: Control saline (F=8; M=7), Complex saline (F=7; M=7); Control amphetamine (F=8; M=8); Complex amphetamine (F=8; M=8).

### Housing procedures

All experimental procedures were carried out following the guidelines set out by the Canadian Council of Animal Care and reviewed by the Animal Welfare Committee at the University of Lethbridge. The control animals were housed in groups of two or three in standard Plexiglas shoebox cages (39 X 57 X 21 cm) with corn-cob bedding covering the floor. The colony room was maintained at 21°C on a 12/12 light dark schedule, with lights on at 7:30 AM.

The complex housing group was lived in large pens measuring 61 x 122 x 183 cm. Three of the walls (sides and front) were made of hardware cloth. The back wall was stainless steel, as was the ceiling and floor. Horizontal platforms were attached to the back wall with runways connecting them. The floor of the structure was covered with corn-cob bedding and various objects were placed on the floor and the wall platforms to encourage exploration. The enclosures were cleaned weekly and new objects introduced each week to maximize exploration. All animals in both conditions had ad libitum access to food and water for the duration of the experiment.

### Amphetamine

At 90 days of age the animals were subdivided into drug and saline treatment conditions. The rats were transported from their housing to a testing room each day for 14 days and placed individually in a Vermax animal activity monitoring system© (open field) for 30 min of habituation prior to drug exposure, during which time locomotion was monitored. Following an injection with d-amphetamine sulfate (1 mg/kg, IP) or saline (0.9%) locomotor activity was monitored for an additional 60 min, after which time the rats were returned to their home cages. The activity monitoring system was equipped with both horizontal and vertical sensors (infra-red beams).

### Anatomy

One week following the last drug or saline treatment the rats were administered an overdose of sodium pentobarbital and perfused intracardially with 0.9% saline. The brains were placed in 20 ml of Golgi-Cox solution and stored in the dark for 14 days and then were placed in a 30% sucrose solution for at least 3 days prior to coronal sectioning (200 um) on a vibrating microtome, mounted on glass slides, and stained using procedures described by Gibb & Kolb (1998). Once the permount had hardened sufficiently the basilar dendrites of layer V pyramidal neurons in the medial prefrontal cortex (Zilles area Cg3) and layer III orbital frontal cortex (Zilles area AID) were quantified, as were the dendrites of medium spiny neurons in the nucleus accumbens shell. Camera lucida was used to draw five neurons from each hemisphere in each area at 250X. Neurons were chosen that were not obscured by other neurons, glia, or blood vessels. Dendritic branching was quantified using the method of Coleman and Reisen (1968).

Dendritic length was estimated using the Sholl (1956) procedure in which an overlay of concentric circles is placed over the neuron drawing beginning at the cell body and spaced 20 microns apart. Spine density was calculated by tracing a length of dendrite (at least 30 um long) at X1000. The exact length of the dendrite was calculated, and the number of spines along the entire length was counted. For cortical pyramidal cells and NAcc cells, spines were counted on one third-order terminal tip per neuron. Spines on the back side of dendrites were obscured so with this method the total spine density counts necessarily underestimate the total spine number. The values for cells in each hemisphere of each rat were averaged, and hemisphere was used as the unit of analysis.

## Results

### Habituation

Locomotor activity during the 30 min habituation period prior to an amphetamine or saline injection decreased over the 14 days of testing, in all groups, as expected. In addition, both the saline and amphetamine control-housed groups show greater habituation than the complex-housed groups (see Figure 1). In addition, after day 1, in which the amphetamine groups received their first amphetamine injection *after* the habituation, the activity of the complex-housed amphetamine group was higher than the complex-housed saline group whereas the control groups did not differ. On average females were more active than males, and the control-housed were more active than the complex-housed groups. Because of the consistent large significant sex difference in activity levels, male and female rats were analyzed separately.

**Figure 1.**
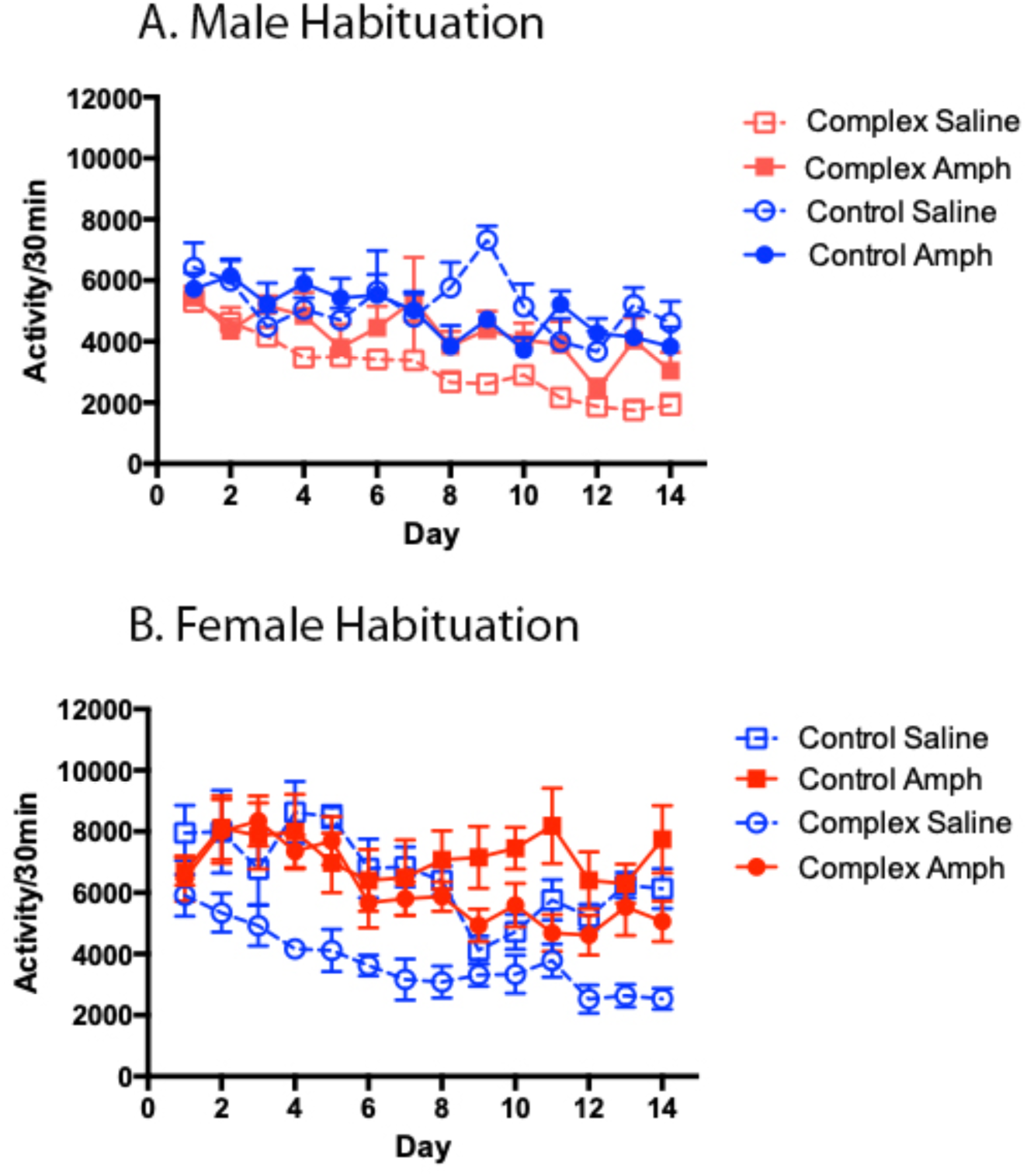
Mean activity during the habituation phase of testing for behavioral sensitization over 14 days. Data = Mean ± SE

Two-way ANOVAs on each sex showed main effects of housing environment (Female; F(1,196)=10.93, p=0.0011), Male; F(13, 140)= 108.3, p<.0001); and test day (Female; F(1,196)=1.752, p=0.0531); Male; F(13, 140)= 2.621, p=.0028)). There was no interaction in the female group (F(13, 196)= 0.9553, p=0.4971) but there was in the male group (F(13, 140)= 2.909, p=0.0009), which reflects no habituation in the male control amphetamine group.

### Acute Amphetamine

Amphetamine significantly increased locomotor activity, relative to saline, in both males and females on Day 1, although females were more active than males, and the effect of amphetamine was attenuated in the complex relative to control group in both sexes (Fig. 2). Two-way ANOVAs showed a main effect of drug (Females F(1,27)=28.44, p<.0001; Males F(1,26)=12.83, p=.0014), and environment for Males F(1,26)=2.786, p=.0379) but not for females F(1,27)=3.283, p<.0811). There was a significant interaction for the Females (F(1,27)=4.249, p=.049) but not for the Males (F(1,26)=1.887, p=.1813). The interaction in the females likely reflects the large activity difference between the control and complex groups. There was a parallel difference in the males, but their activity levels were much lower than those for the females. In any event the data show that housing in the complex environment attenuated the acute locomotor activating effect of amphetamine relative to the controls.

**Figure 2.**
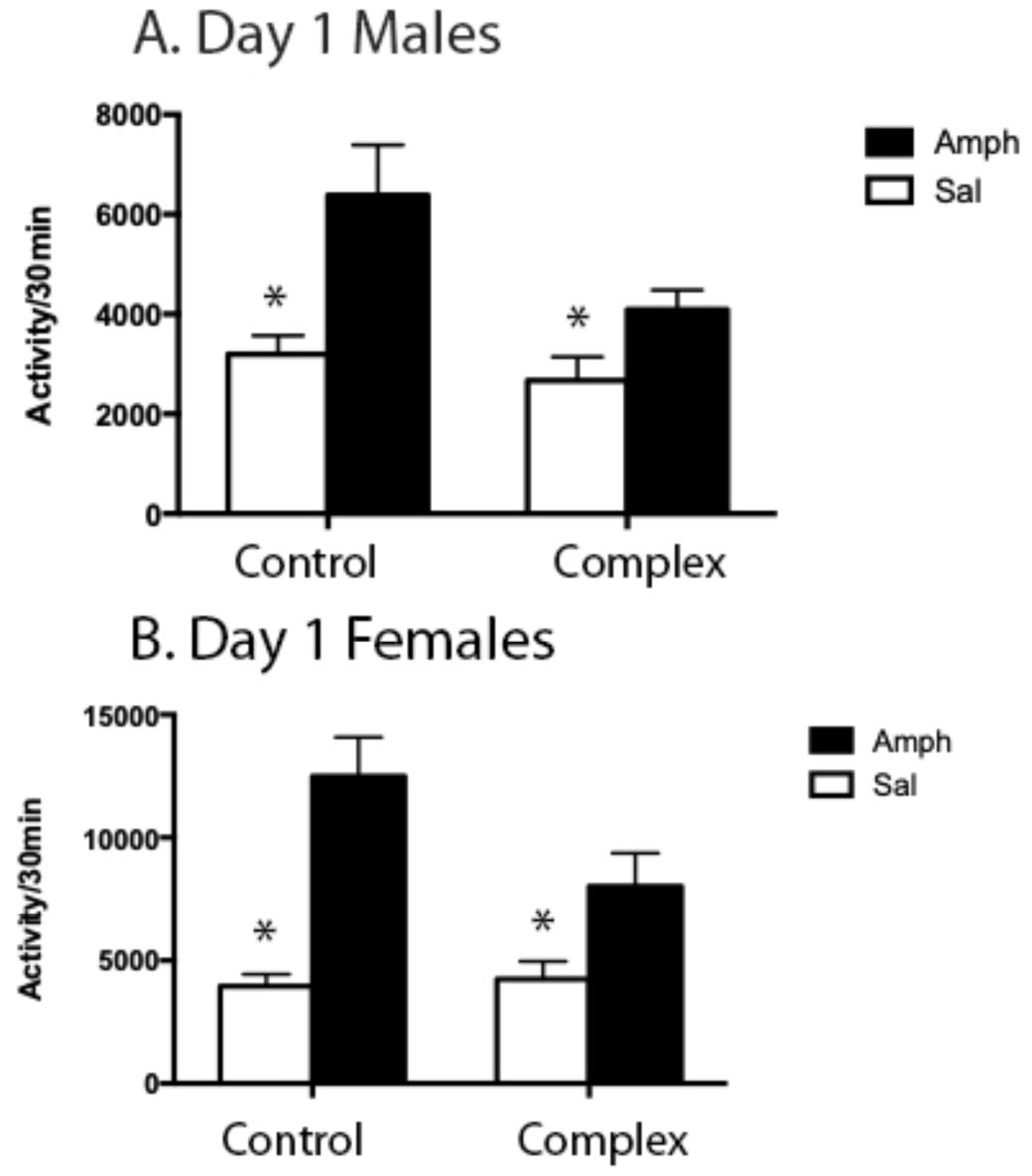
Comparison of the effect of amphetamine on control-housed and complex-housed animals on day 1 of amphetamine administration. All groups showed increased activity in the amphetamine-treated animals. Data = Mean ± SE. *p<.05 or better

### Repeated Amphetamine

Amphetamine-induced locomotor activity increased across the 14 days of treatment in all groups; that is, repeated daily amphetamine treatment produced behavioral sensitization in both complex and control-housed males and females (Fig. 3). Females were more active than males on both the first and last day of treatment, but there was no sex difference in the magnitude of the change in locomotor activity across time, suggesting no sex difference in the degree of behavioral sensitization. However, females did sensitize more rapidly and plateau sooner (with fewer injections) than males. The complex-housed group was less active than the control-housed group on both the first and last day of testing, such that there was no effect of housing condition on the degree of change across time (i.e., in sensitization), in either males or females.

**Figure 3.**
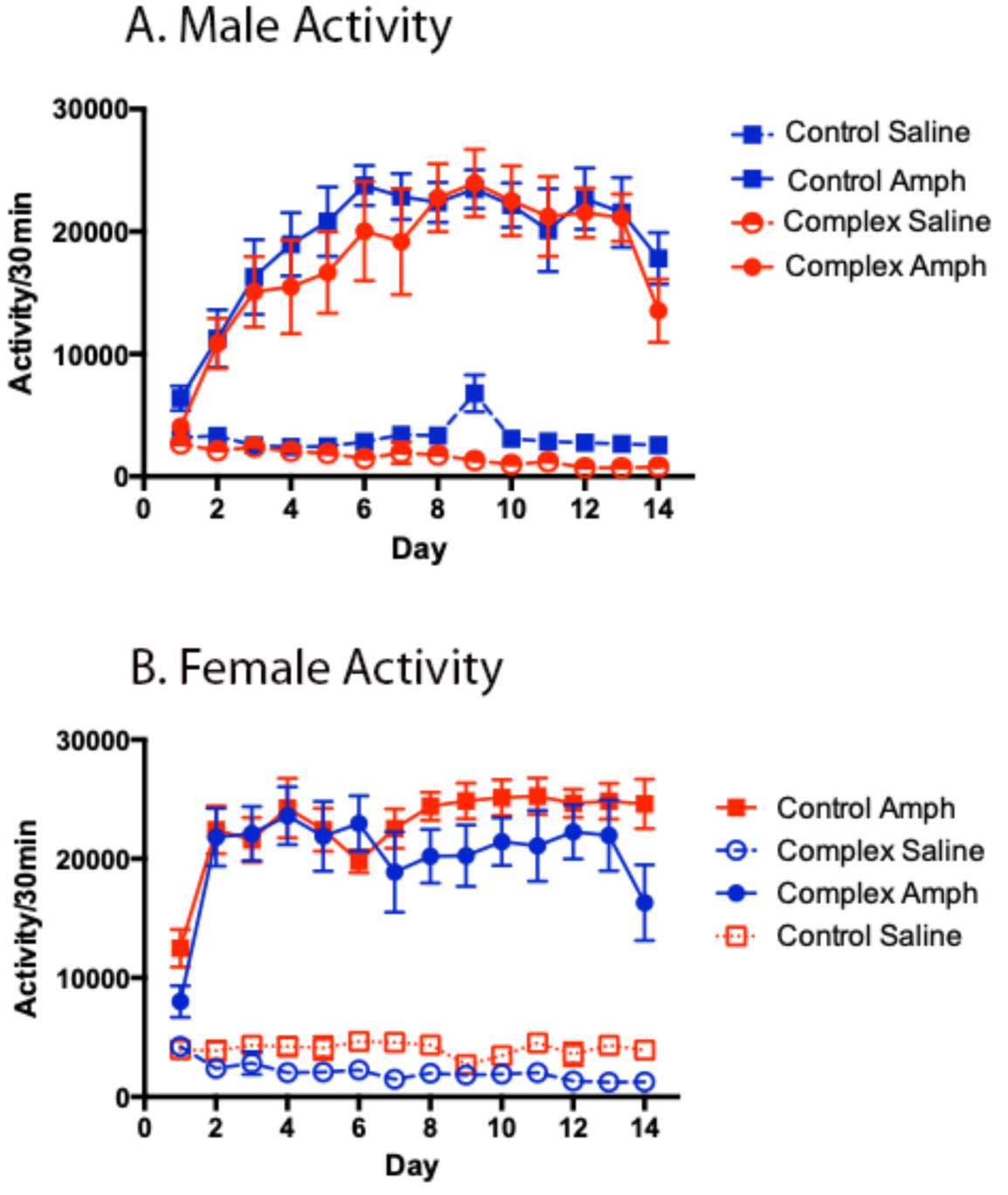
Mean activity during the amphetamine-administration phase of testing for behavioral sensitization over 14 days. Data = Mean ± SE

ANOVAs comparing day 1 to day 14 activity shows significant effects for day (Female: (F(1,50)=11.91, p=.0011) (Male: (F, 1,49)=19.66, p<.0001), environment (Female: (F(1,50)=36.09, p<.0001) (Male: (F, 3,49)=23.62, p<.0001), and the interactions (Female: (F(1,50)=8.173, p=.0002) (Male: (F, 3,49)=11.21, p<.0001. The interactions reflect the fact that the activity went up in the amphetamine groups and down in the saline groups.

#### Anatomy

Three different questions were asked related to dendritic anatomy: 1) Were there effects of housing condition (enrichment)? 2) Were there effects of amphetamine treatment? And, 3) were there any housing by amphetamine interactions? In addition, there is the more general question of whether there are sex differences in any of the above. To simplify the presentation of the data, we will consider each question separately but also discuss the effect of sex in each case. Two-way ANOVAs were done, with housing condition and drug treatment as effects. We note that the degrees of freedom may vary slightly across regions owing to differences in acceptable staining. The staining was generally good and comparable to our previous studies (e.g., Robinson & Kolb, 1997). Again, because there were obvious sex differences in some anatomical measures, we analyzed the sexes separately, in part to avoid complex three-way interactions.

#### Effect of housing condition in males?

There was an effect of housing condition in all three regions, but it varied by region (see Figures 4-6). Thus, in Cg3 there was a *decrease* in dendritic length and branching but an *increase* in spine density in rats housed in the complex environment. In AID there were no changes in dendritic length or branching but a *decrease* in spine density.

**Figure 4.**
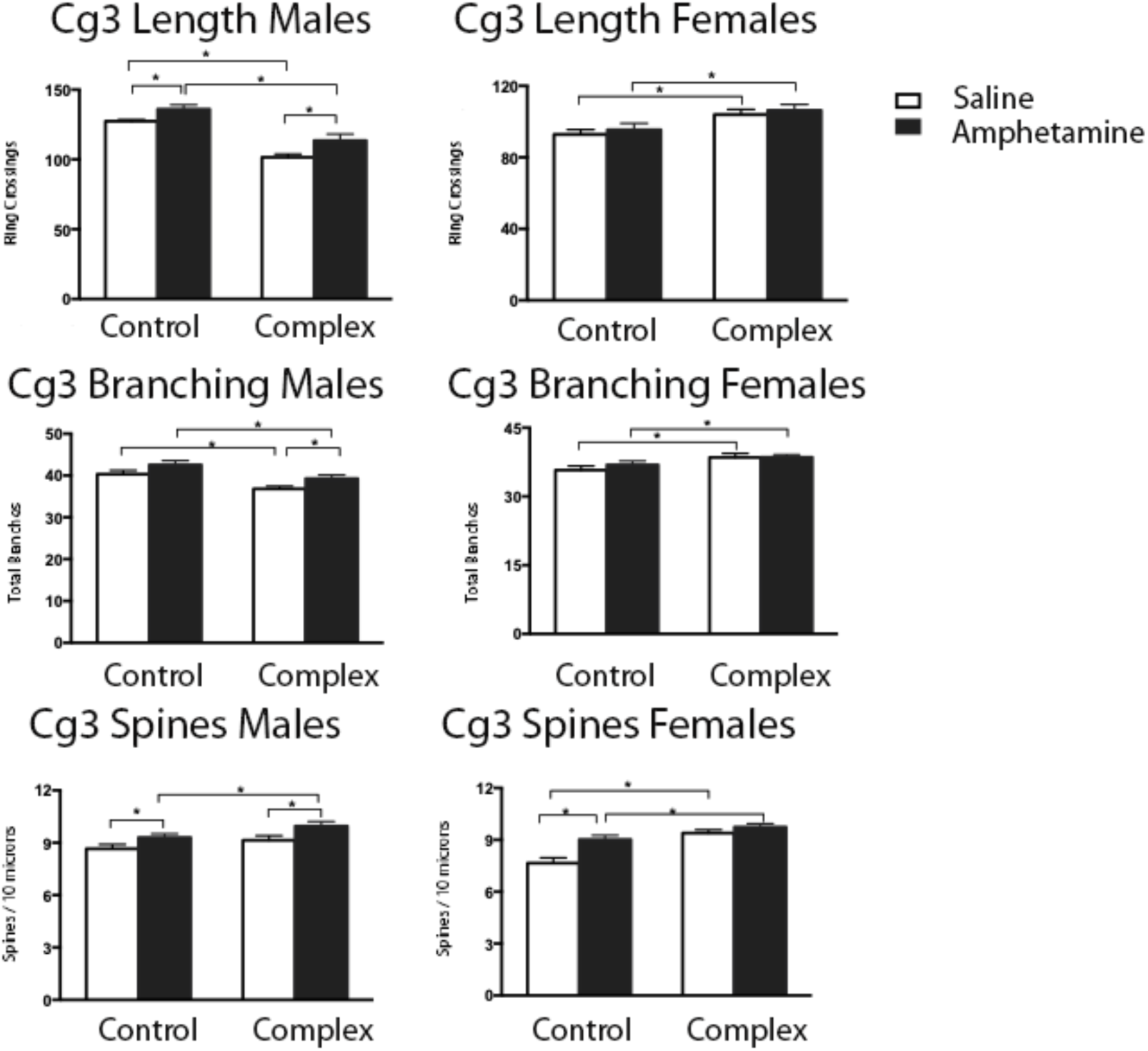
Summary of the effects of amphetamine and complex housing on dendritic measures in Zilles’ area Cg3. Data = Mean ± SE. p<.05 or better

**Figure 5.**
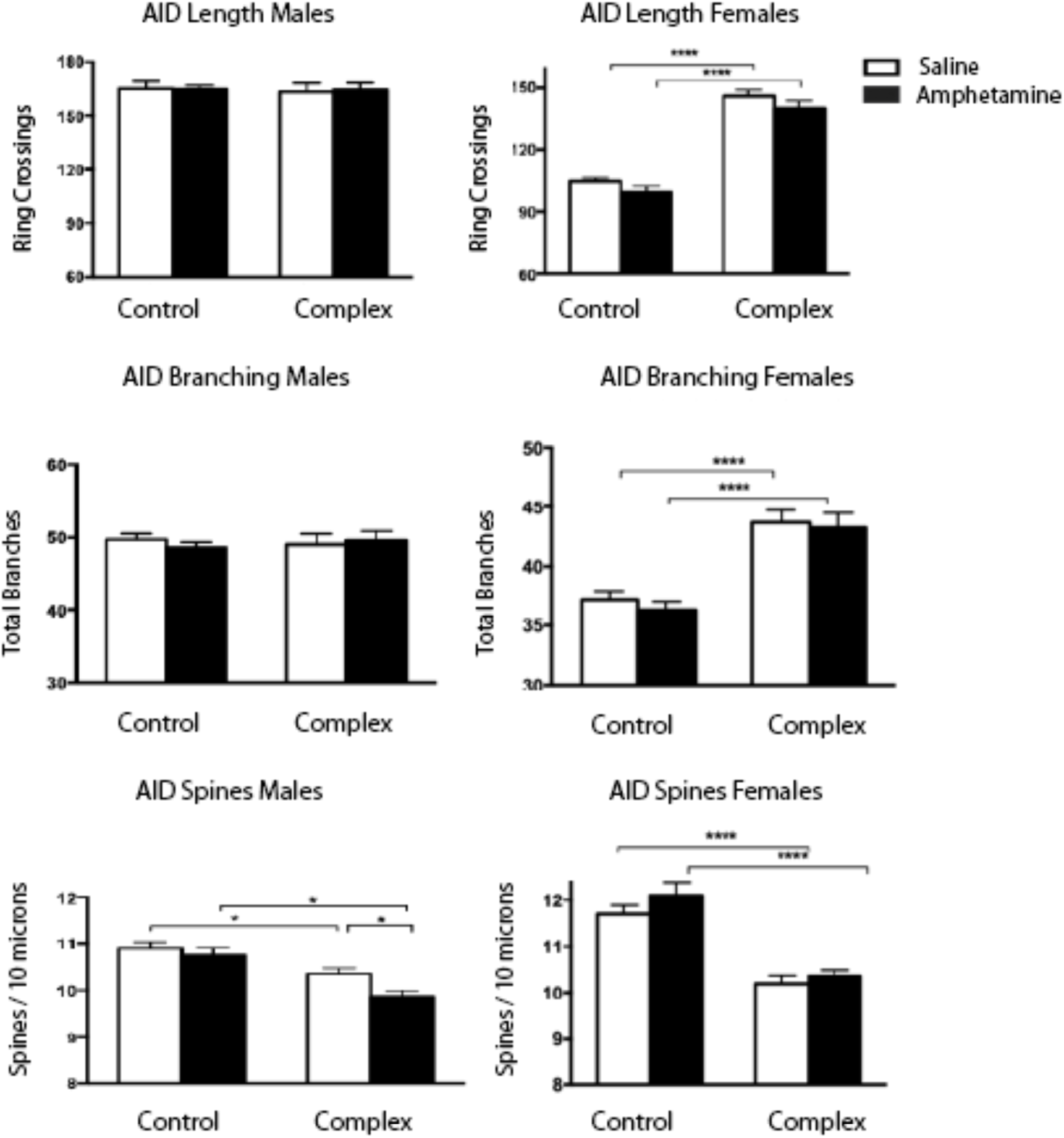
Summary of the effects of amphetamine and complex housing on dendritic measures in Zilles’ area AID. Data = Mean ± SE. p<.05 or better

**Figure 6.**
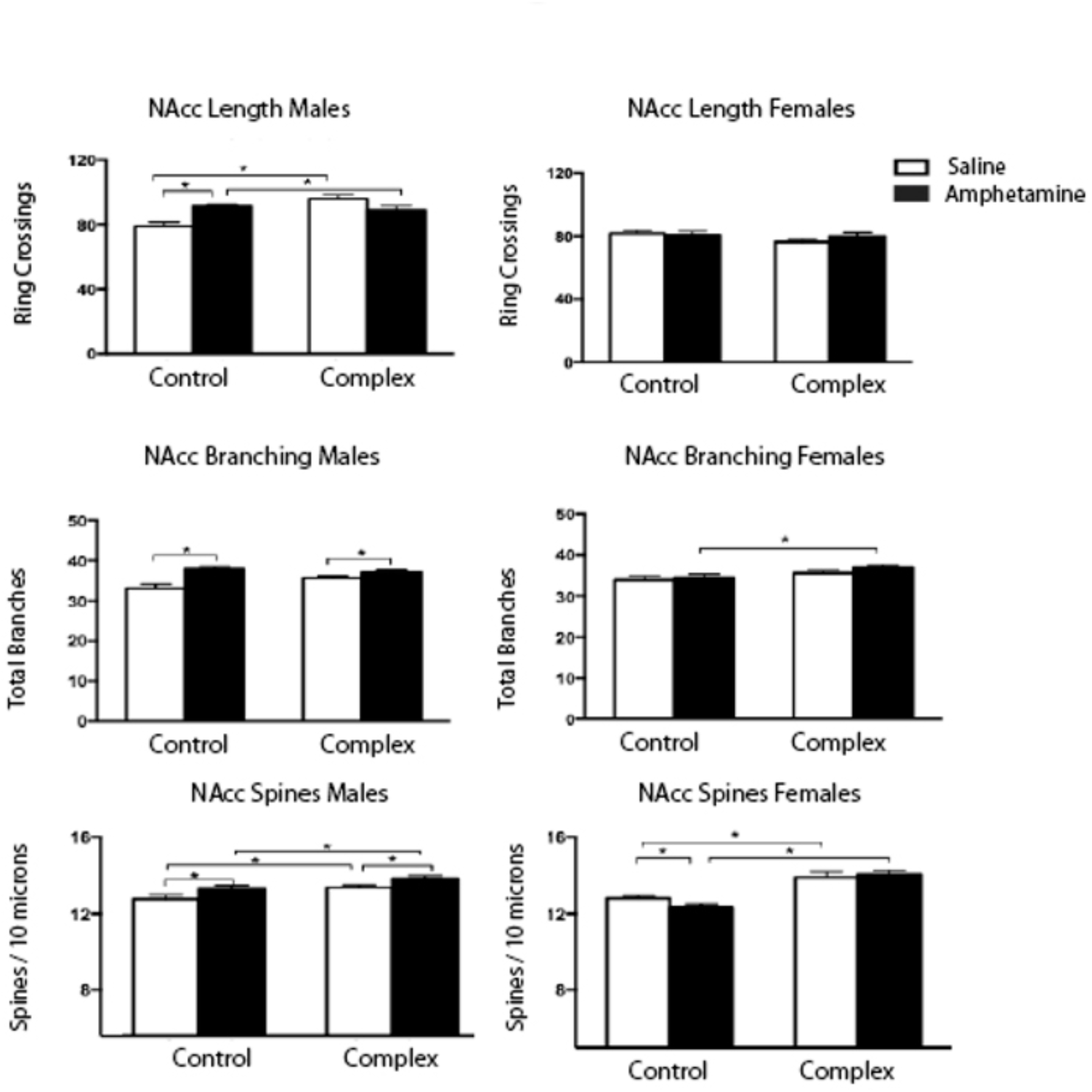
Summary of the effects of amphetamine and complex housing on dendritic measures in NAc. Data = Mean ± SE. p<.05 or better

In NAcc, complex housing was associated with an increase in length and spine density. ANOVA for Cg3 showed a significant effect of complex housing for length (F(1,50)=51.67, p<.0001), branching (F(1,54)=15.61, p=.0002), and spine density (F(1,58)=5.334, p<.0245). ANOVA for AID showed no effect for length (F(1,58) = 0.06, p=.8023) or branching (F(1,58)=0.017, p=.8954), but there was a significant decrease in spine density (F(1,58)=32.91, p<.0001).

ANOVA for NAcc found a significant increase in dendritic length (F(1,48)=7.856, p=.0073), spine density (F(1,52)=10.35, p<.0001), but not for branching (F(1,50)=2.664, p=.1089).

#### Effect of housing condition in females?

In sharp contrast to the males, there were increases in all measures except dendritic length in NAcc, where there were no effects (see Fig 6). ANOVA for Cg 3 showed a significant housing effect showing *increased* length (F(1,52)=12.35, p=.0009), branching (F(1,54)=7.495, p=.002), and spine density (F(1,50)=27.82, p<.0001). ANOVA for AID showed a significant *increase* in dendritic length (F(1,49)=197.4, p<.0001) and branching (F(1,49)=55.06, p<.0001), but a *decrease* spine density (F(1,52)=71.03, p<.0001). ANOVA for NAcc showed a significant *increase* in branching (F(1,58)=6.438, p=.0139), and spine density (F(1,52) =39.19, p<.0001), but no effect on length (F(1,54)=2.182, p=.1454). Thus, not only was there a large enrichment effect in the females, overall, the effect was large.

We note, too, that the dendritic length and branching were larger in male than female controls, confirming our previous findings in control animals (Kolb & Stewart, 1991).

#### Effect of amphetamine in males?

Amphetamine *increased* dendritic length and spine density in both Cg3 and NAcc and branching in the NAcc but not Cg3. There was no effect of amphetamine on length or branching in AID, but a *decrease* in spine density. ANOVA for Cg3 showed a significant enrichment effect for *increased* length (F(1,50)=9.402, p=.0035), branching (F(1,54)=7.44, p=.0086), and spine density (F(1,58)=8.713, p=.0046). ANOVA for AID showed no effect for length (F(1,58) = 0.001, p=.969) nor branching (F(1,58)=0.034, p=.8539), but there was a significant *decrease* in spine density (F(1,58)=6.2434, p<.0155). ANOVA for NAcc found a significant *increase* in dendritic branching (F(1,50 )=26.83, p=.0001), and spine density (F(1,52)=8.876, p=.0048), but not for length (F(1,48)=1.342, p=.2524).

#### Effect of amphetamine in females?

In contrast to the males, there were virtually no drug effects in the females. The only significant effects were an *increase* in spine density Cg3 and a *decrease* in spine density in NAcc. ANOVA for Cg3 showed no main effect for length nor branching (F(1,52)=0.623, p=.4336), ((F(1,52)=0.534, p=.4681), but there was for spine density (F(1,50)=13.31, p=.0006). ANOVA for AID found no main effect of any measure (F(1,49)=0.4823, p=.4906), (F(1,49)=3.750, p=.0586), (F(1,52)=2.014, p=0.1619), respectively.

ANOVA for NAcc found no main effect of any measure (F(1,58)=1.247, p=.2687), (F(1,54)=0.639, p=.5489), (F(1,52)=.4939, p=0.4853), respectively.

#### Interaction effects?

There were virtually no housing condition X amphetamine interactions. The only significant effects were for dendritic branching and length in male NAcc (1,48)=15.26, p=.0003), (F(1,50)=7.684, p=.0078) and female Cg3 spine density (1,50)=17.453, p=.0342). The male NAcc effects reflect an amphetamine effect in controls but not complex-housed animals. The Cg3 female spine interaction again reflected an amphetamine effect in control but not complex housed animals.

Taken together, the anatomical results show a greater effect of housing condition than amphetamine overall and that complex housing did not influence the effects of amphetamine, which confirms our earlier study in which nicotine administration was also unaffected by complex housing. The small amphetamine effect in females was unexpected.

## Discussion

There are four major findings. 1) Relative to the control group, both male and female complex-housed animals showed greater habituation to the drug test environment over days of testing. 2) Both standard and complex-housed males and females showed a similar degree of locomotor sensitization as a function of repeated treatment with amphetamine, but females sensitized more rapidly than males and the complex-housed animals (both males and females) were generally less active. 3) There were large effects of housing condition on dendritic anatomy, but these varied markedly by anatomical region (Cg3, AID, NAcc) and by sex. 4) Treatment with amphetamine also produced large effects on dendritic measures in both Cg3 and NAcc and spine measures in all three regions of male rats. There were no effects on dendritic measures in female and amphetamine affected spine density only in Cg3 (*increased* density) and NAcc (*decreased* density) (see Table 1). We will discuss each finding in turn before considering possible explanations for the effects of housing condition on both behavior and brain.

**Table 1.**
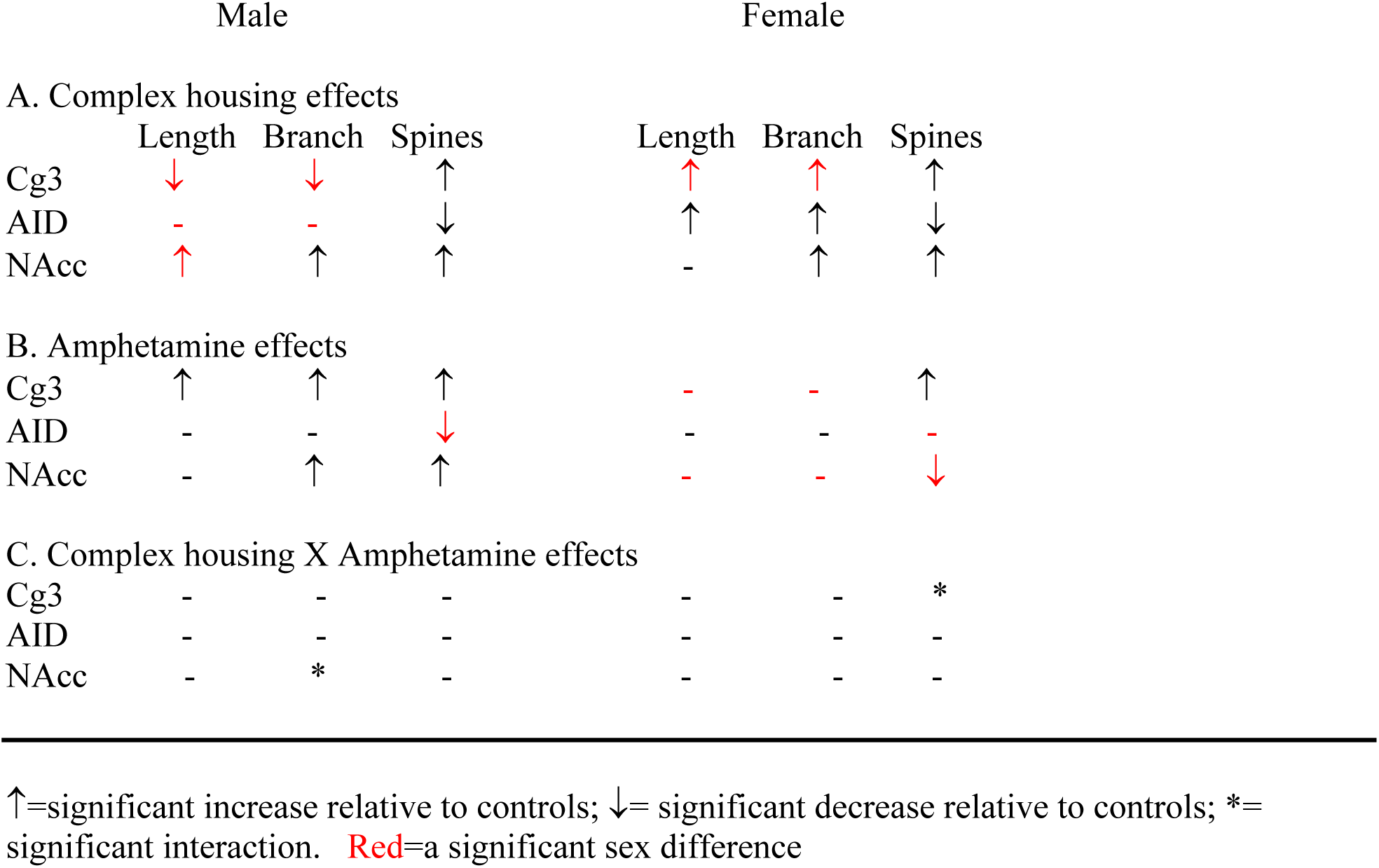
Summary of anatomical results

### Locomotor habituation prior to drug administration

Relative to the control-housed groups the complex-housed groups showed greater habituation in locomotor activity seen when they were initially placed in the test chamber, prior to drug administration, across the 14 days of testing. This could reflect the fact that the complex-housed groups would typically have more spontaneous activity in their usual environment and, given that their environment changed often, they had more experience with novelty.

### Effects of amphetamine on locomotor activity

As expected, the first injection of amphetamine increased locomotor activity in all groups, and both sexes, but importantly, there was a clear effect of housing condition. Acute amphetamine produced a much smaller increase in locomotor activity in animals raised in the complex environment. It is not clear what that difference reflects, but as in the habituation phase, the difference could be related to the fact that the complex-housed animals were accustomed to a much larger and stimulating environment. However, after the initial injection of amphetamine, both control and complex housed male and female animals developed locomotor sensitization to a comparable degree, indicating the housing condition had a negligible effect on this form of experience-dependent plasticity. This may be related to the fact that complex housing also did not prevent most effects of amphetamine on dendritic morphology. However, females did sensitize more rapidly than males, consistent with previous reports of sex differences in psychomotor sensitization (Robinson 1984; Becker et al., 2006 for review).

### Regional and sex differences in the effects of complex housing on neuronal morphology

We are unaware of any previous studies on the effects of complex housing from conception to adulthood in lab animals. A study by DeRosa et al (2022) found that housing dams and pups in complex environments had a significant effect on constituents of maternal milk, which underscores the multidimensional impact of maternal and neonatal environments on brain and behavioral development in the offspring. One effect was a change in the epigenetic profile of the milk. We have shown previously that the same dose of amphetamine as in the current paper changed the pattern of gene expression in mPFC, orbitofrontal cortex (including AID), and NAcc in male Long-Evans rats, although the pattern of changes was different in each region (Mychasiuk et al., 2012). (Females were not studied.) We can now make three predictions.

First, if the mother’s milk has a different pattern of gene expression in complex vs control environments, we can predict that there could be a different pattern of gene expression in brain regions in the offspring. Second, given that complex housing also changes gene expression in brain (Rampon et al., 2000), we can predict that the effects of amphetamine administration in animals whose gene expression has already been altered by complex housing will likely be different than gene expression seen in control animals. Third, given that experiences can produce a different pattern of gene expression in males and females (Mychasiuk et al., 2012), we can predict that complex housing will produce a different a different pattern of gene expression in females than males. Thus, given that different patterns of gene expression in cortex would likely produce different patterns of neuronal morphology, we could expect a different pattern of neuronal organization in the two sexes, which we found in the current study.

We had shown in a previous study (Kolb et al., 2003c) that housing females in complex environments for 3.5 mo, beginning at 110 days of age, had no effect on dendritic length or branching in mPFC, although it did increase those measures in parietal cortex and NAcc. (Males were not studied, nor was the AID.) In the current study in which complex housing began at conception, there were increases in dendritic length and branching in females but a decrease on the same measures in males. Thus, it appears that complex housing alters dendritic morphology (length and branching) in the mPFC when animals are housed there from conception, but not when placed in a complex environment beginning in adulthood, perhaps due to the kinds of factors discussed by DeRosa et al (2022).

### Regional and sex differences in the effects of amphetamine on neuronal morphology

The sex differences in the amphetamine effects on neuronal morphology were unexpected given that the behavioral effects were comparable in the two sexes, and we have consistently seen similar amphetamine-related changes in neuronal morphology in Cg1 and NAcc in female and male rats (e.g., Muhammad & Kolb, B., 2011). There is, however, convincing evidence for sex differences in the effects of methamphetamine on brain and behavior in rats, and structural brain changes seen in noninvasive imaging in humans (Daiwile et al., 2022). Furthermore, the social effects of drug use are different in men and women (e.g., Hser et al., 2005; Lin et al., 2004).

Differences in social effects of drug treatment may be important in the current study. One significant difference in the methods between our earlier studies and the current one is that in our earlier studies the animals were housed singly during the amphetamine administration period, whereas in the current study the control-housed animals were group housed and following the amphetamine activity testing the animals were returned to their respective group housing. Access to a social partner is reinforcing and has positive incentive value (e.g., Achterberg et al., 2016; Vanderschuren et al., 2016) and increases in extracellular dopamine increases the positive reinforcing effects of social contact in both male and female rats (Sharp & Smith, 2022). The details of social interactions are reported to influence DNA methylation in both prefrontal cortex and hippocampus (Mychasiuk et al., 2011), which in turn is likely to alter neuronal morphology.

Thus, social interactions post drug exposure might significantly alter the effects of psychoactive drugs in a sexually dimorphic manner, as seen in the current study. It is possible that group housing, even in the control-housed animals, was a sufficiently complex environment to occlude some effects of amphetamine in females, but not males. The fact that complex housing changed neuronal morphology in females in the current study would rule out the possibility that the female brain is less plastic than male brains, but given the sex difference in the enrichment effects on neuronal morphology observed here, it appears that female and male brains are plastic in different ways (see also Daiwile et al., 2022).

Taken together, the anatomical results show a greater effect of housing environment than amphetamine overall. The small amphetamine effect in females was unexpected and in contrast to our earlier studies, but as noted above this may be related to the fact that in the current study the control-housed animals were group housed. We should also note that the post-drug survival times were longer in our previous studies, which could have affected the results. There was also large sex difference of enrichment effects, especially in AID where there were large effects in dendritic length and branching in females, but no effect on either measure in males.

## Conclusion

The effects of psychomotor stimulant drugs on brain morphology are clearly influenced by a lifetime of experience. Given that humans typically have a lifetime of experiences before being exposed to psychoactive drugs, it seems likely that the anatomical effects of drug taking are somewhat different than one might predict from our earlier studies using rats with more limited experiences. Researchers designing future studies of drug effects on brain and behavior may be well advised to keep these findings in mind (see also Kentner et al., 2021).

